# Reduced transmission of *Mycobacterium africanum* compared to *M. tuberculosis* in urban West Africa

**DOI:** 10.1101/206136

**Authors:** Prince Asare, Adwoa Asante-Poku, Diana Ahu Prah, Sonia Borrell, Stephen Osei-Wusu, Isaac Darko Otchere, Audrey Forson, Gloria Adjapong, Kwadwo Ansah Koram, Sebastien Gagneux, Dorothy Yeboah-Manu

## Abstract

**Background:** Understanding transmission dynamics is useful for tuberculosis (TB) control. We conducted a population-based molecular epidemiological study to understand TB transmission in Ghana.

**Methods:** *Mycobacterium tuberculosis* complex (MTBC) isolates obtained from prospectively-sampled pulmonary TB patients between July, 2012 and December, 2015 were confirmed as MTBC using IS*6110* PCR. MTBC lineages were identified by large sequence polymorphism and single nucleotide polymorphism assays and further characterized using spoligotyping and standard 15-loci MIRU-VNTR typing. We used the n-1 method to estimate recent TB transmission and identified associated risk factors using logistic regression analysis.

**Findings:** Out of 2,309 MTBC isolates, we identified 1,082 (46·9%) single cases with 1,227 (53·1%) isolates belonging to one of 276 clustered cases (clustering range; 2-35). Recent TB transmission rate was estimated to be 41·2%. While we see no significant difference in the recent transmission rates between lineages of *Mycobacterium africanum* (lineage-5 (31·8%); lineage-6 (24·7%), p=0·118), we found that lineage-4 belonging to the *M. tuberculosis* transmitted significantly higher (44·9%, p<0·001). Finally, apart from age being significantly associated with recent TB transmission (p=0·007), we additionally identified a significant departure in the male/female ratio among very large clustered cases compared to the general TB patient population (3:1 vs. 2:1, p=0·022).

**Interpretations:** Our findings indicate high recent TB transmission suggesting occurrences of unsuspected outbreaks. The observed reduced transmission rate of *M. africanum* suggests other factor(s) may be responsible for its continuous presence in West Africa.

**Funding:** Wellcome Trust Intermediate Fellowship Grant 097134/Z/11/Z to Dorothy Yeboah-Manu.

## Research in context

### Evidence before this study

We searched PubMed for population-based molecular epidemiology studies conducted in Ghana before May, 2011 and again June 2017 using the search terms “tuberculosis” and “transmission”, “population based”, “population-based” or “epidemiology”. We found no such study conducted in Ghana and confirmed this with coordinators of the National Tuberculosis control Program (NTP). Till date, the only detailed population-based molecular epidemiological study conducted in Ghana has been from our research group where we found the distribution of *Mycobacterium africanum* and *Mycobacterium tuberculosis* sensu stricto to be fairly constant in the respective proportions of 20% and 80% within the study period. However, to the best of our knowledge, no study has been conducted to identify the transmission dynamics of circulating MTBC strains in Ghana and this makes this current study the first of its kind.

### Added value of the study

This study, for the first time in Ghana using combined discriminatory power of spoligotyping and 15-loci MIRU-VNTR typing, has estimated the recent TB transmission rate (clustering rate) to be 41·2% with the urban and rural settings having estimated rates of 41·7% and 9·0% respectively. Within our study population, we identified the unlikeliness of a clustered case to harbor either isoniazid or rifampicin resistant TB strain (adjusted OR 0·7, 95% CI 0·5 – 0·9) and showed low transmission of multidrug resistant TB strains. From this study, we have also identified that each year increase in age is significantly associated with an approximately 1% (CI: 0·13 – 2·00, p=0·007) decrease in the odds of a TB patient being part of a recent transmission event putting age as a significant risk factor. This implies that, compared to younger individuals, older individuals are more likely to get active TB disease by reactivation of latent TB infection. Also, we identified that the male/female ratio among very large clusters (cluster size; n>20) was significantly higher than that observed in the general TB patient population (3:1 vs. 2:1, p=0·022) with some large clusters (cluster size between 6 and 20) involving only males. Using adjusted predictions, we found that TB patients were more likely to be involved in a recent transmission event in 2012 compared to 2013 – 2015 and further identified that, clustering generally increases with increase in the number of TB contacts. The low observed transmission among Maf lineages and their constant prevalence as previously reported indicate other factors(s) (environmental/host) is (are) likely to be responsible for maintaining Maf in Ghana.

### Implication of all the available evidence

The slight reduction of transmission rate after 2012 might be as a result of increased awareness during our national tuberculosis survey conducted in 2013. This shows that, with increased awareness we can reduce TB burden resulting from recent transmission. Despite the numerous efforts by the NTP, in 2015 Ghana recorded a very low TB detection rate of 34% (14,999/44,000) which is way below the African and global targets of 50% and 70% respectively. The rate of sputum conversion (having microscopy results showing negative for AFB among TB patients) and relapse following the intensive 2 months, at 5 months or the entire 6 months TB treatment regimen has been used by the NTP as conventional indicators for assessing TB control. Some aspects of TB control such as identifying the frequency of reactivation of latent infection and the risk of recent TB transmission among the population would be missed by the currently used conventional indicators. Our current study provides vital information which will help the control program.

## Introduction

Tuberculosis (TB) is a global health emergency; in 2015 an estimated 10.4 million people got sick, while 1.8 million died of TB.^1^ The World Health Organization (WHO) in 1993 declared TB a global health emergency and called for much efforts and resources to fight this global menace. Due to the largely inefficacy of the Bacillus Calmette-Guérin (BCG) vaccine, the current TB control strategy especially in resource limited settings relies on case detection and treatment by the Directly Observed Therapy short course (DOTs) strategy. The conventional indicators used for assessing the national control programs under this strategy focus on the proportion of patients with new sputum smear positive pulmonary disease that are cured by the end of treatment or whose sputum microscopy becomes negative after the first 2 months of treatment. Such indicators ignore equally important aspects of TB control, which includes the duration of infectivity, the frequency of reactivation, and the risk of progression among the infected contacts and the proportion of TB due to recent transmission.

Understanding transmission dynamics will contribute to knowledge on factors that enhance spread of the disease, which is useful for TB control. Molecular epidemiological studies have been very useful in a number of countries in supporting TB control by identifying populations at most risk, transmission sites as well as providing much understanding on the prevalence of different *Mycobacterium tuberculosis* complex (MTBC) strains with varied virulence and drug resistance incidence rates.^2–5^ These studies have shown that the dynamics of TB transmission vary greatly geographically. Even though Africa harbors a large proportion of the global TB cases with current incidence at 275 per 100,000 population,^1^ population-based molecular epidemiological studies needed to understand transmission patterns are rare, with the few studies conducted lacking in-depth analysis of the transmission dynamics of MTBC strains belonging to different lineages.^6–8^

The molecular typing tools, spacer oligonucleotide typing (spoligotyping) and mycobacterial interspersed repetitive unit-variable number of tandem repeat (MIRU-VNTR) typing, have been successfully used for strain differentiation in TB transmission studies due to their combined high discriminatory power and reproducibility, and in combination with epidemiological data, have been remarkably used for the detection of recent TB transmission/outbreaks.^2,4,9,10^ Currently, due to the high cost and expertise needed for employing whole genome sequencing and analysis, population-based studies may not readily employ its use and considering capacity building in a low resource setting like Ghana, spoligotyping and MIRU-VNTR typing remains good alternatives.

Tuberculosis in humans is caused mainly by *Mycobacterium tuberculosis* sensu stricto (MTBss) and *M. africanum* (Maf) which are further divided into seven lineages [MTBss: lineage 1-4 and 7 (L1-L4 and L7), Maf: lineages 5 and 6 (L5 and L6)]^11,12^; while MTBss is globally distributed 13 Maf is restricted to West Africa where it is responsible for up to 50% of TB cases.^13^ Nevertheless, reports mainly from the Gambia where L6 is prevalent suggest Maf is attenuated compared to MTBss given the indication that Maf will be outcompeted by MTBss.^11,14,15^ However, an 8-year study recently conducted in Ghana found the prevalence of Maf to be fairly constant with approximately 20% indicating that Maf may transmit as equally as MTBss (mainly L4).^16^ The objective of the study was to determine the transmission dynamics of TB caused by MTBss and Maf.

## Methods

### Study design and population

This study was a population-based prospective study in which sputum samples were collected from consecutive clinically diagnosed pulmonary TB patients reporting to 12 selected health facilities within an urban setting [Accra Metropolitan Assembly (AMA)] and a rural setting East Mamprusi district (MamE) (Figure 1). The study was conducted from July, 2012 to December, 2015. A pulmonary TB case was defined as an individual with both clinical and bacteriologically confirmed case of TB. Detailed demographic and epidemiological data including age, sex, employment status, residential address, health status and previous history of TB were obtained from each participant after obtaining consent using a structured questionnaire.

**Figure 1:**
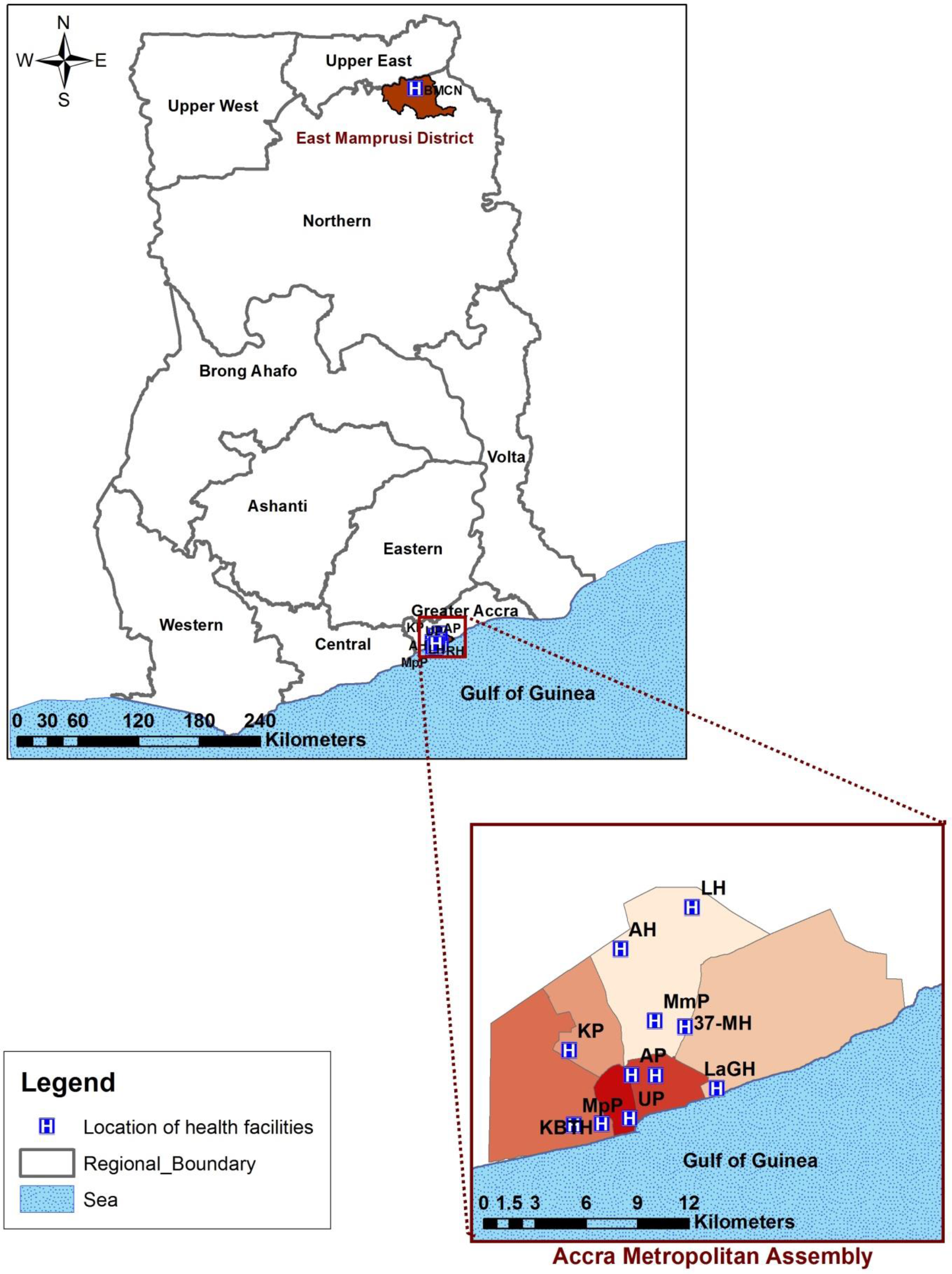
Map of study area showing the location of the 12 sampling sites (health facilities)

### Mycobacterial isolation, species identification and drug susceptibility testing

The sputum samples were decontaminated and cultured on Lowenstein-Jensen media to obtain mycobacterial isolates which were confirmed as MTBC by detecting the MTBC-specific 17 insertion sequence IS*6110* using PCR.^17^ *In vitro* drug susceptibility to isoniazid and rifampicin were determined using either the microplate alamar blue cell viability assay as described elsewhere,^18^ and/or the Geno-type MTBDR*plus* (Hain lifescience), according to the manufacturer’s protocol.^19^

### Lineage and strain classification

Lineage and strain classification of the MTBC was achieved in a step wise manner using large sequence polymorphism typing identifying regions of difference 4, 9, 12, 702, and 711, ^11,13^ single nucleotide polymorphism typing, spoligotyping,^20^ and MIRU-VNTR typing.^21^ For MIRU-VNTR typing, we first used a customized set of 8-MIRU loci as described by Asante-Poku *et al.*^22^ and resolved clustered cases by analyzing the remaining 7 loci of the standard MIRU-15 loci set.^21^ All assays were well controlled with PCR amplifications and pre-PCR procedures conducted in physically separated compartments to avoid laboratory cross contamination. The presence of more than one allelic repeat number (multiple allele) for any given locus is suggestive of laboratory cross contamination, multiple strain infection or microevolution of a single strain. To prevent bias resulting from cross contamination and multiple strain infection, isolates with multiple allele at more than one MIRU loci (described as *untypeable*) were excluded from further analysis. Isolates with only one multiple allele at any given locus were however included due to the possibility of microevolution.

The spoligotyping patterns and assigned shared type numbers obtained were defined according to SITVITWEB database (http://www.pasteur-guadeloupe.fr:8081/SITVIT_0NLINE/) while sublineages were assigned based on the MIRU-VNTRplus database (http://www.miru-vntrplus.org).^23^ Strains with no lineage nomenclature data were further identified using the TB-lineage database^24^ or otherwise regarded as orphan strains. A strain was defined as an MTBC isolate with a unique molecular signature thus, a unique spoligotype pattern and/or a unique MIRU-VNTR allelic pattern for the number of investigated MIRU loci.

### Clustering analysis and risk factor assessment

Clustering analysis was performed using the categorical parameter and the UPGMA coefficient from a constructed phylogenetic tree using the online MIRU-VNTR tool. Clustering analysis was based on the assumption that, strains with the same DNA fingerprint may be epidemiologically linked and associated with recent TB transmission.^25^ A Cluster was defined as two or more isolates (same strain) that share an indistinguishable spoligotype and 15-loci MIRU-VNTR allelic pattern but allowing for one missing allelic data at any one of the *difficult-to-amplify* MIRU loci (VNTR 2163, 3690 and 4156). We also defined the size of a cluster using the total number of isolates in the cluster into categories of small (2 isolates), medium (3–5 isolates), large (6–20 isolates) and, very large (>20 isolates).

The recent transmission rate was estimated using the n-1 formulae; 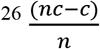

Where; *nc* is the total number of clustered cases, *c* is the number of clusters, and *n* is the total number of cases in the sample.

We included only one strain per participant in our analysis and excluded follow-up cases. Clustering analysis was stratified first by location and then by MTBC lineage. The spatial distribution and clustering among all of the observed Spoligo/MIRU strain types were studied by constructing a minimum spanning tree (MST) with Bionumerics software (Applied Maths, Sint-Marteen-Latem, Belgium).

### Data management and analysis

Both molecular and epidemiological data were analyzed. Epidemiological data retrieved from all participants with positive MTBC cultures were included in the analysis while excluding data from those with no growth, contaminated cultures and isolated non-tuberculous mycobacterial species. All statistical analyses were carried out using the Stata statistical package version 14·2 (Stata Corp., College Station, TX, USA). The association of specific lineages and/or sub-lineages of the MTBC with time and/or geographical locations were explored using chi-square test and a logistic regression model. For the determination of independent predictive factors for recent TB transmission, a multivariate analysis (forward step-wise approach with a probability entry of 0·1) was conducted using a logistic regression model while estimating the odds ratios (OR). P-values < 0·05 were considered significant.

The manuscript was reported according to the “strengthening the reporting of molecular epidemiology for infectious diseases (STROME-ID)” guidelines.^27^

### Ethics statement

Ethical clearance for this study was obtained from the Scientific and Technical Committee and then the Institutional Review Board at NMIMR, University of Ghana with a federal wide assurance number FWA00001824 after reviewing and approving the protocols and procedures.

## Results

### Characteristics of study participants

We recruited 3,303 sputum smear positive pulmonary TB cases and obtained 2,604 (78·8%) MTBC isolates. After excluding 13 *M. bovis* and isolates that were untypeable (described in methods), 2,309/2,604 (88·7%) isolates were included for clustering analysis. The participants included comprised 71% (1,631) males and 29% (663) females (15 participants had no record of gender) with a median age of 39 (range: 3 to 91) and 33 (range: 4 to 90), respectively (Figure 2; appendix p 2). The male/female ratio observed was comparable to the national average of approximately 2:1. Of the 2,309 MTBC participants, 201 (8·7%) were from the rural setting and 2,108 (91·3%) from the urban setting. Among our study cohort, 7·4% (184/2,482) of participants were previously treated cases, which is similar to the national value of 7·2%^1^. Seventy-one percent (1,561/2,208) presented with a bacterial burden resulting from sputum smear microscopy of at least 2+ and 33% (544/1,665) admitted having contact with at least one TB patient. In a multivariate logistic regression analysis, we found that male patients are less likely to be infected with a L5 strain (adjusted OR 0·7, 95% CI 0·5 – 0·9) and individuals living in villages are more likely to be infected with a L6 strain (OR 6·6, CI 1·2 – 36·1) (appendix p 3).

**Figure 2:**
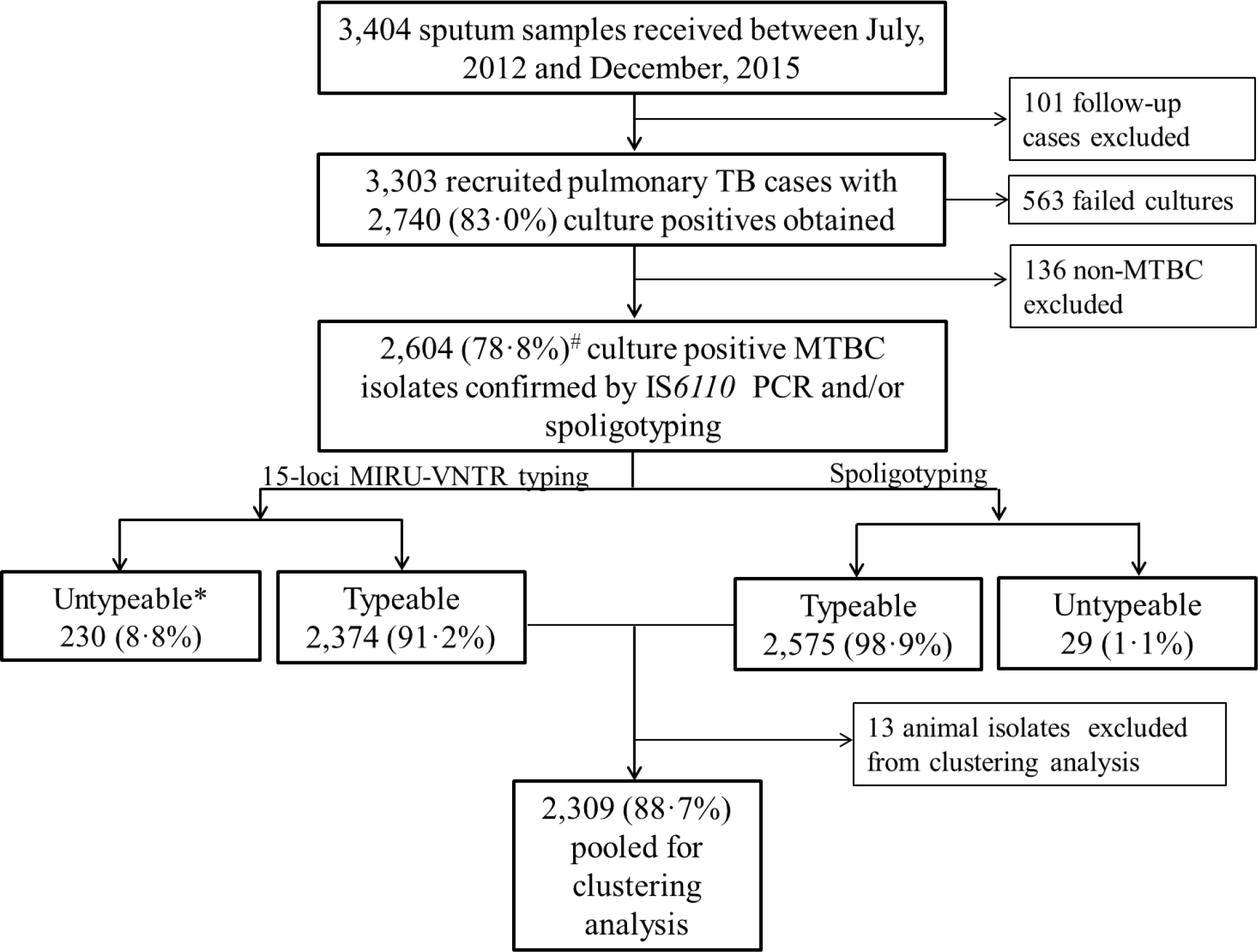
Pipeline for recruited participants and culture positive TB cases used for clustering analysis. *Category described as untypeable for MIRU-VNTR includes isolates with ≥ 2 MIRU-loci un-amplified (164, 71·3%) and isolates with double allele at ≥ 2 MIRU-loci (66, 28·7%). These isolates were described as suspected mix infection or laboratory contamination and hence were removed from further analysis. # Frequency was expressed as the total number of *Mycobacterium tuberculosis* complex (MTBC) isolates obtained

### Population structure and recent transmission rate estimation

Among the 2,309 MTBC isolates analyzed for clustering, 1,870 (81·0%) were MTBss and 439 (19·0%) were Maf. Six of the seven human-adapted MTBC lineages were found, with L4, L5 and L6 being most frequent with 1,741 (75·4%), 289 (12·5%), and 150 (6·5%) isolates, respectively (table 1). The relative proportions of the most frequent MTBC lineages remained constant over the entire three and half year study period (ptrend: L4 p=0·7168, L5 p=0·8379, L6 p=0·2466, Figure 3).

**Figure 3:**
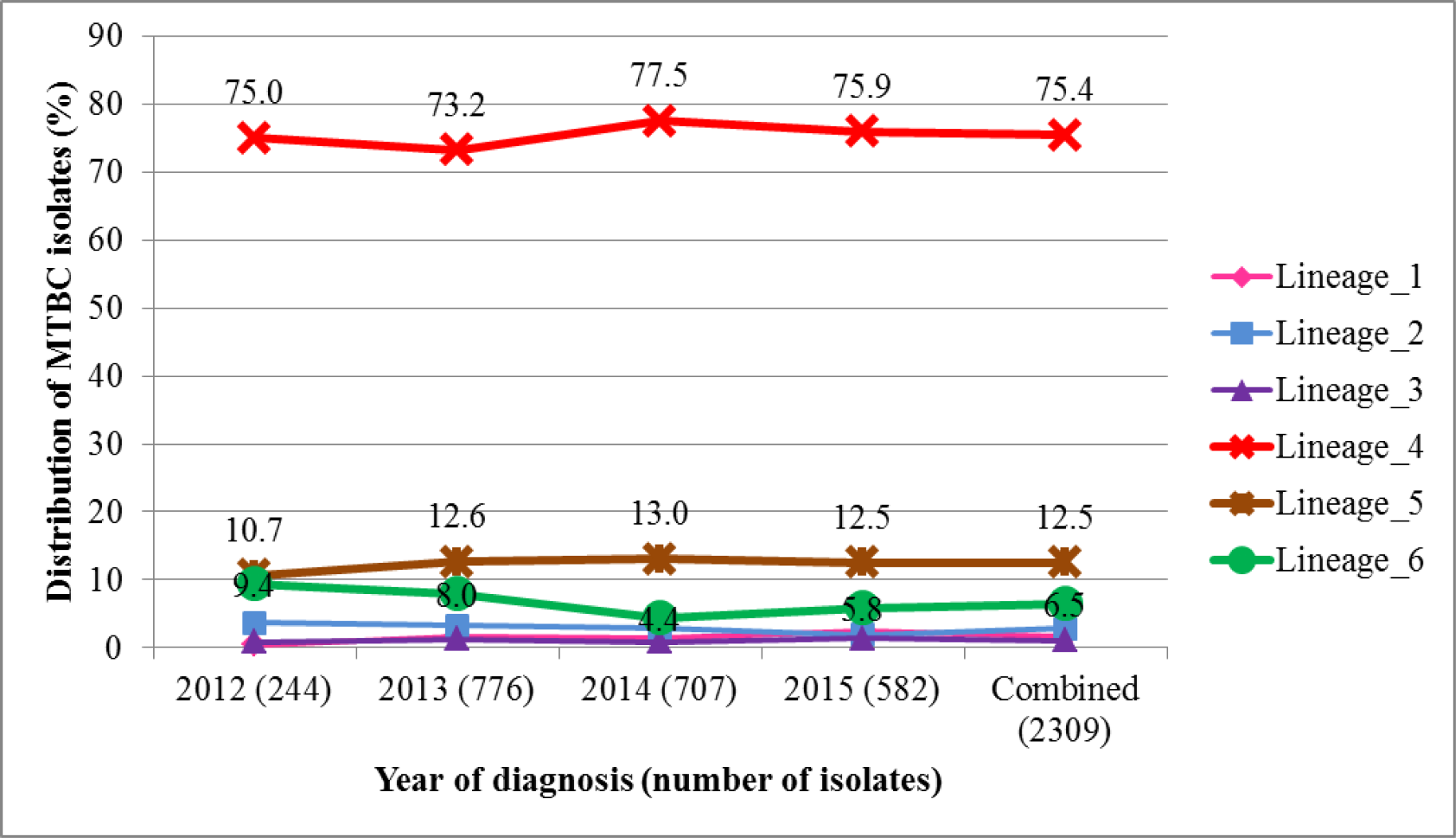
Temporal distribution of 2,309 MTBC isolates stratified by lineage. Lineages have been color coded with the universally accepted color codes for the main MTBC lineages.

**Table 1:**
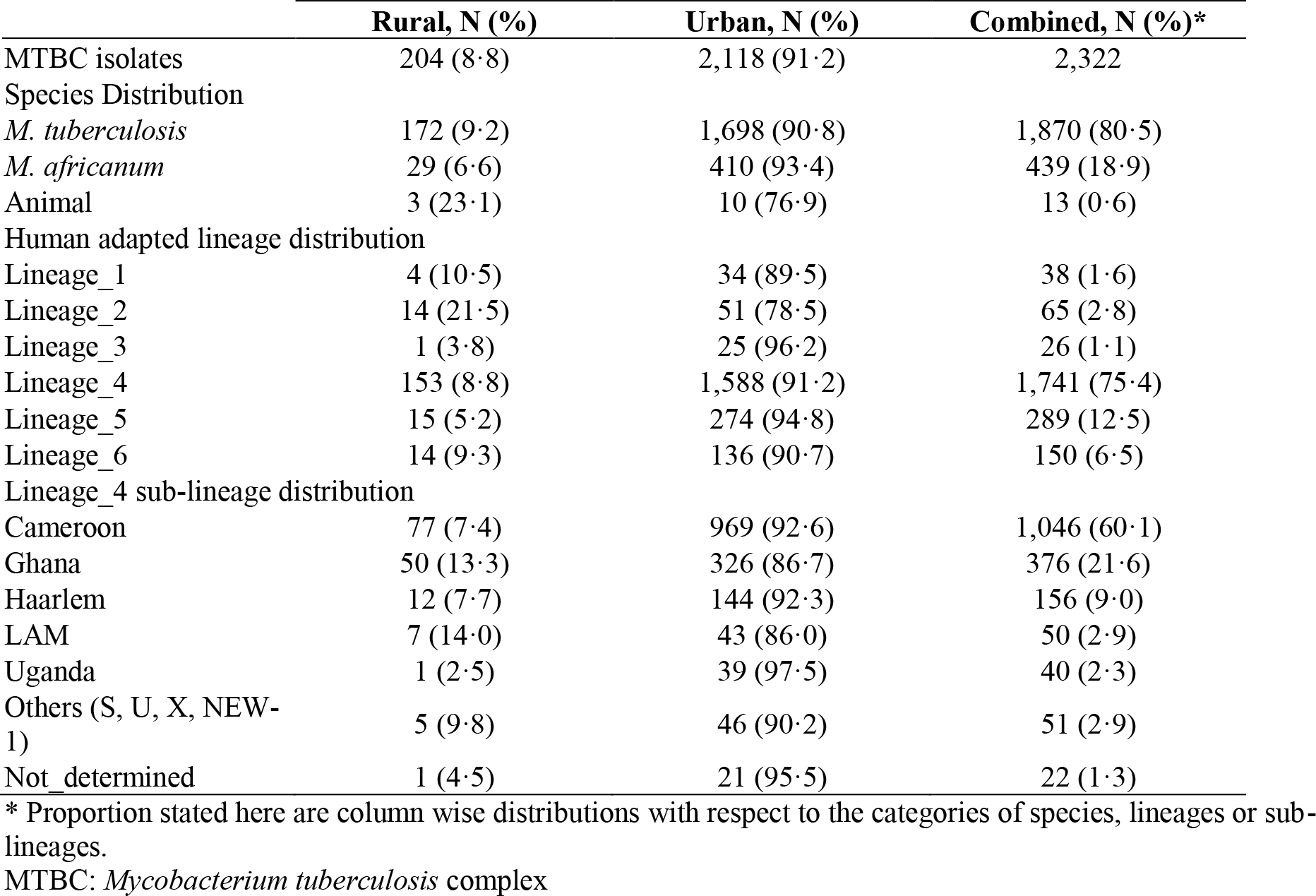
Geographical distribution and population structure of MTBC in Ghana by spoligotyping

|  | Rural, N (%) | Urban, N (%) | Combined, N (%)* |
| --- | --- | --- | --- |
| MTBC isolates | 204 (8.8) | 2,118 (91.2) | 2,322 |
| Species Distribution |  |  |  |
| *M. tuberculosis* | 172 (9.2) | 1,698 (90.8) | 1,870 (80.5) |
| *M. africanum* | 29 (6.6) | 410 (93.4) | 439 (18.9) |
| Animal | 3 (23.1) | 10 (76.9) | 13 (0.6) |
| Human adapted lineage distribution |  |  |  |
| Lineage_1 | 4 (10.5) | 34 (89.5) | 38 (1.6) |
| Lineage_2 | 14 (21.5) | 51 (78.5) | 65 (2.8) |
| Lineage_3 | 1 (3.8) | 25 (96.2) | 26 (1.1) |
| Lineage_4 | 153 (8.8) | 1,588 (91.2) | 1,741 (75.4) |
| Lineage_5 | 15 (5.2) | 274 (94.8) | 289 (12.5) |
| Lineage_6 | 14 (9.3) | 136 (90.7) | 150 (6.5) |
| Lineage_4 sub-lineage distribution |  |  |  |
| Cameroon | 77 (7.4) | 969 (92.6) | 1,046 (60.1) |
| Ghana | 50 (13.3) | 326 (86.7) | 376 (21.6) |
| Haarlem | 12 (7.7) | 144 (92.3) | 156 (9.0) |
| LAM | 7 (14.0) | 43 (86.0) | 50 (2.9) |
| Uganda | 1 (2.5) | 39 (97.5) | 40 (2.3) |
| Others (S, U, X, NEW-1) | 5 (9.8) | 46 (90.2) | 51 (2.9) |
| Not_determined | 1 (4.5) | 21 (95.5) | 22 (1.3) |

\* Proportion stated here are column wise distributions with respect to the categories of species, lineages or sub-lineages.
MTBC: *Mycobacterium tuberculosis* complex

Of the 2,309 isolates included for clustering analysis, we identified 1,227 (53·1%) isolates being clustered in 276 different clusters with an average cluster size of 4 (range: 2 − 35) and 1,082 (46·9%) singletons, giving a total of at least 1,358 unique MTBC strains circulating within our study population (table 2a). Using the n-1 method, we estimated the overall clustering rate (reflecting recent transmission rate) to be 41·2%. Lineages 2, 4 and 5 contributed high clustering rates of 53·8%, 44·9%, and 31·8%, respectively (table 2a). The Cameroon, Ghana and Haarlem sub-lineages of L4 were the most abundant sub-lineages and significantly contributed to the observed high L4 clustering rate (p<0·05) and there was no significant difference between the Cameroon and Ghana sub-lineages (p=0·565) (Figure 4). While we see no significant difference in the recent transmission rates between members of Maf (L5 and L6, p=0·118), we found that L4 transmitted significantly higher (p<0·001) with seven of its clusters having very large cluster sizes (>20 isolates per cluster) made up of the Ghana sub-lineage (4 very large clusters) and Cameroon sub-lineage (3 very large clusters) (Figure 4; appendix p 6). Notwithstanding the lower transmissibility of lineages 5 and 6, we also observed four large clusters for each of these lineages. The urban and rural settings had estimated recent transmission rates of 41·7% and 9·0% respectively.

**Figure 4:**
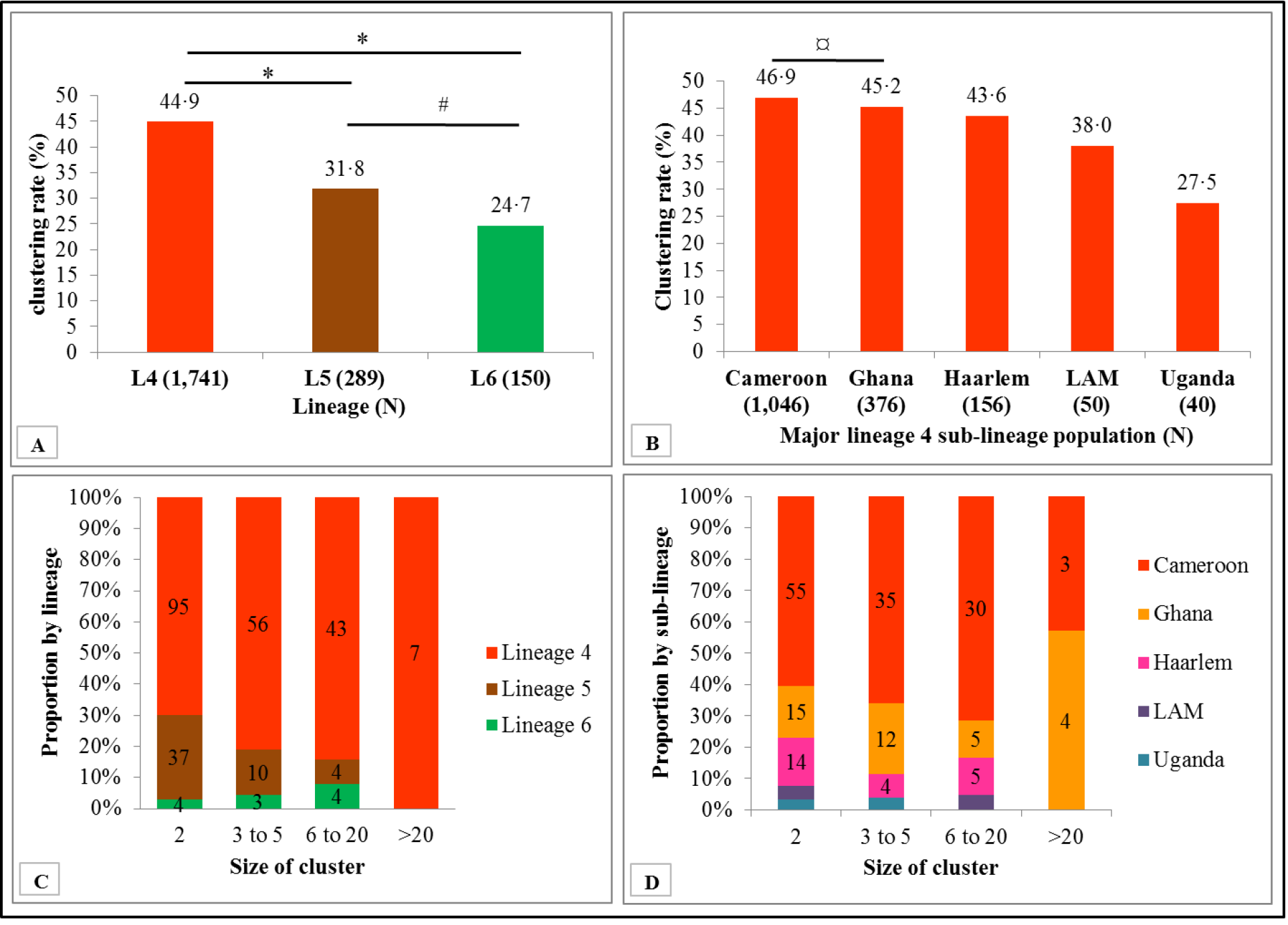
Cluster distribution and size stratified by lineage (panel A and C) and sub-lineage (panel B and D). *p<0·001 #p=0·118 ¤p=0·565

**Table 2a:** Clustering analysis stratified by lineages and major sub-lineage populations of MTBC

| Lineage | Isolates (n) | Clustered cases (c) | Clustered strains (nc) | Single cases (s) | Total strain types (s+c) | Clustering rate<sup>†</sup> (%) |
| --- | --- | --- | --- | --- | --- | --- |
| Lineage 1 | 38 | 3 | 7 | 31 | 34 | 10.5 |
| Lineage 2 | 65 | 8 | 43 | 22 | 30 | 53.8 |
| Lineage 3 | 26 | 2 | 4 | 22 | 24 | 7.7 |
| Lineage 4 | 1,741 | 201 | 982 | 759 | 960 | 44.9 |
| Cameroon<sup>#</sup> | 1046 | 123 | 614 | 432 | 555 | 46.9 |
| Ghana<sup>#</sup> | 376 | 36 | 206 | 170 | 206 | 45.2 |
| Haarlem<sup>#</sup> | 156 | 23 | 91 | 65 | 88 | 43.6 |
| LAM<sup>#</sup> | 50 | 6 | 25 | 25 | 31 | 38.0 |
| Uganda<sup>#</sup> | 40 | 5 | 16 | 24 | 29 | 27.5 |
| Lineage 5 | 289 | 51 | 143 | 146 | 197 | 31.8 |
| Lineage 6 | 150 | 11 | 48 | 102 | 113 | 24.7 |
| Summary* | 2309 | 276 | 1227 | 1082 | 1358 | 41.2 |

\* Summary was calculated using only the items in cells corresponding to the six main lineages.
<sup>#</sup> Major lineage 4 sub-population
<sup>†</sup> The clustering rate was used for estimating the recent transmission rate

**Table 2b:** Clustering analysis stratified by study setting and lineages/major sub-lineage populations of MTBC

| Lineage | Isolates (n) |  | Clustered cases (c) |  | Clustered strains (nc) |  | Single cases (s) |  | Total strain types (s+c) |  | Clustering rate<sup>†</sup> (%) |  |
| --- | --- | --- | --- | --- | --- | --- | --- | --- | --- | --- | --- | --- |
|  | Urban | Rural | Urban | Rural | Urban | Rural | Urban | Rural | Urban | Rural | Urban | Rural |
| Lineage 1 | 34 | 4 | 3 | 0 | 7 | 0 | 27 | 4 | 30 | 4 | 11.8 | 0 |
| Lineage 2 | 51 | 14 | 5 | 1 | 33 | 4 | 18 | 10 | 23 | 11 | 54.9 | 21.4 |
| Lineage 3 | 25 | 1 | 2 | 0 | 4 | 0 | 21 | 1 | 23 | 1 | 8 | 0 |
| Lineage 4 | 1,588 | 153 | 183 | 10 | 907 | 25 | 681 | 128 | 864 | 138 | 45.6 | 9.8 |
| Cameroon<sup>#</sup> | 969 | 77 | 112 | 5 | 575 | 10 | 394 | 67 | 506 | 72 | 47.8 | 6.5 |
| Ghana<sup>#</sup> | 326 | 50 | 32 | 4 | 182 | 12 | 144 | 38 | 176 | 42 | 46 | 16 |
| Haarlem<sup>#</sup> | 144 | 12 | 20 | 1 | 81 | 3 | 63 | 9 | 83 | 10 | 42.4 | 16.7 |
| LAM<sup>#</sup> | 43 | 7 | 6 | 0 | 25 | 0 | 18 | 7 | 24 | 7 | 44.2 | 0 |
| Uganda<sup>#</sup> | 39 | 1 | 5 | 0 | 16 | 0 | 23 | 1 | 28 | 1 | 28.2 | 0 |
| Lineage 5 | 274 | 15 | 49 | 0 | 137 | 0 | 137 | 15 | 186 | 15 | 32.1 | 0 |
| Lineage 6 | 136 | 14 | 10 | 0 | 43 | 0 | 93 | 14 | 103 | 14 | 24.3 | 0 |
| Summary | 2108 | 201 | 252 | 11 | 1131 | 29 | 977 | 172 | 1229 | 183 | 41.7 | 9 |

\* Summary was calculated using only the items in cells corresponding to the six main lineages.
<sup>#</sup> Major lineage 4 sub-population
† The clustering rate was used for estimating the recent transmission rate

### Exploring the diversity and clustering within the MTBC lineages

We included data from both 15-MIRU (allelic information) and spoligotyping pattern to construct minimum spanning trees. Very large molecular clusters (clusters with > 20 isolates: *defined in methods*) were observed for L4 in addition to one strikingly large cluster belonging to the Beijing family of lineage 2 (Figure 5; appendix p 7). Generally, we observed low distribution of multidrug resistant TB strains across all the major lineages (appendix p 8 − 10). There was no single large cluster with all isolates being multidrug resistant with only rare focal MDR-TB distribution among some lineage 4 clusters (appendix p 8). The spatial distributions of the isolates constituting each cluster stratified by study setting are shown in appendix (p 11 – 13).

**Figure 5:**
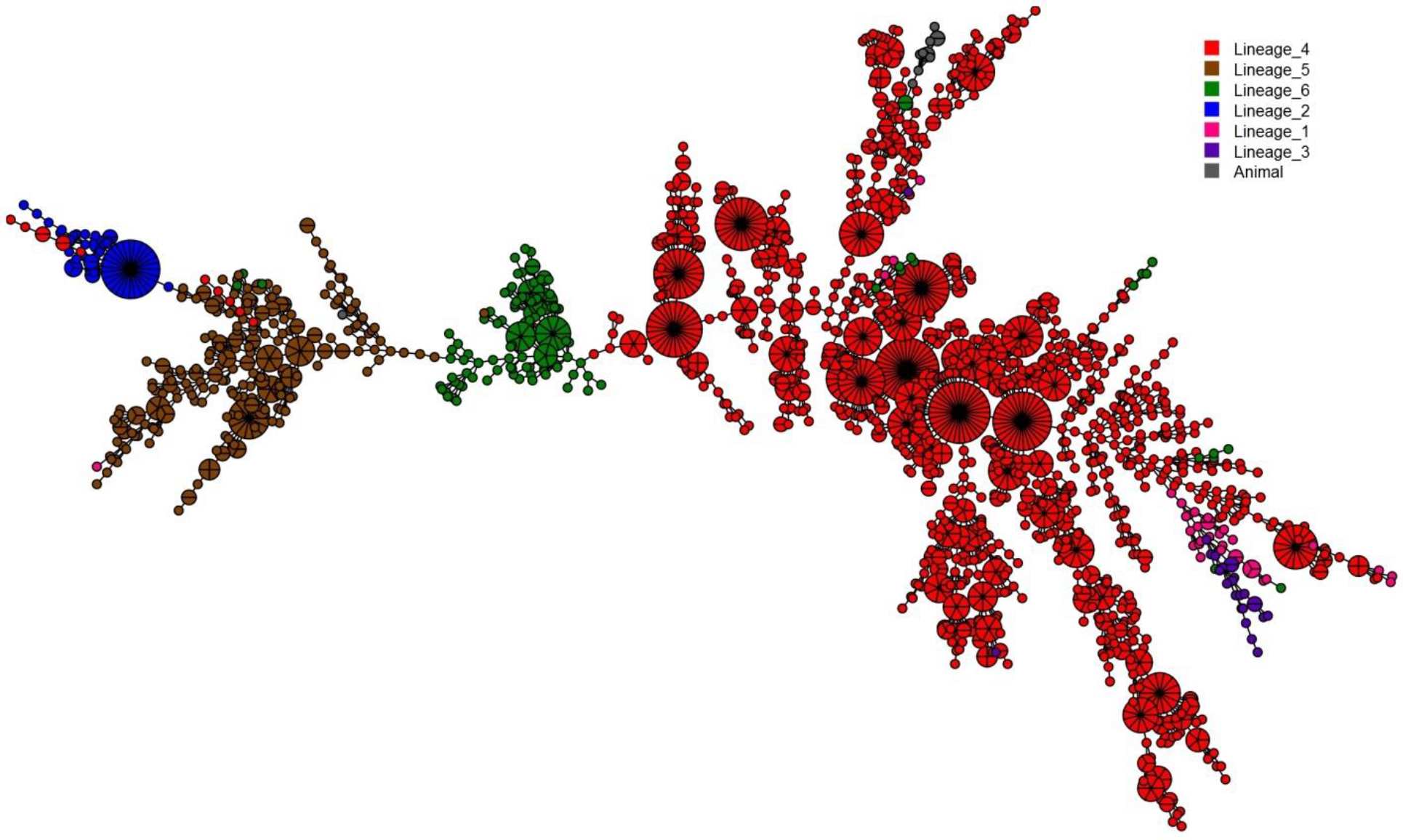
Minimum spanning tree (MST) representation of the clustering of 2,322 MTBC isolates from Accra Metropolitan Assembly and East Mamprusi district built with Bionumerics software. The color code reflects the main MTBC lineages 1 to 6 with the size depicting the number of clustered isolates with an identical strain type.

### Molecular epidemiology and factors associated with clustering: Logistic regression modeling

Next we looked for risk factors that may associate with recent TB transmission. We identified a total of 675 individuals belonging to either large (6 − 20 isolates) or very large (>20 isolates) molecular clusters with a combined median cluster size of 14 (range 6 − 35). Majority of the individuals belonging to very large clusters were males with a male to female ratio of approximately 3:1, significantly higher than the 2:1 ratio observed in the general TB patient population (p=0·022). Three large clusters; cluster ID MSC4193, MSC5003 X and MSC4107 with cluster sizes of 9, 7 and 7 respectively, involved only males (table 3).

**Table 3:**
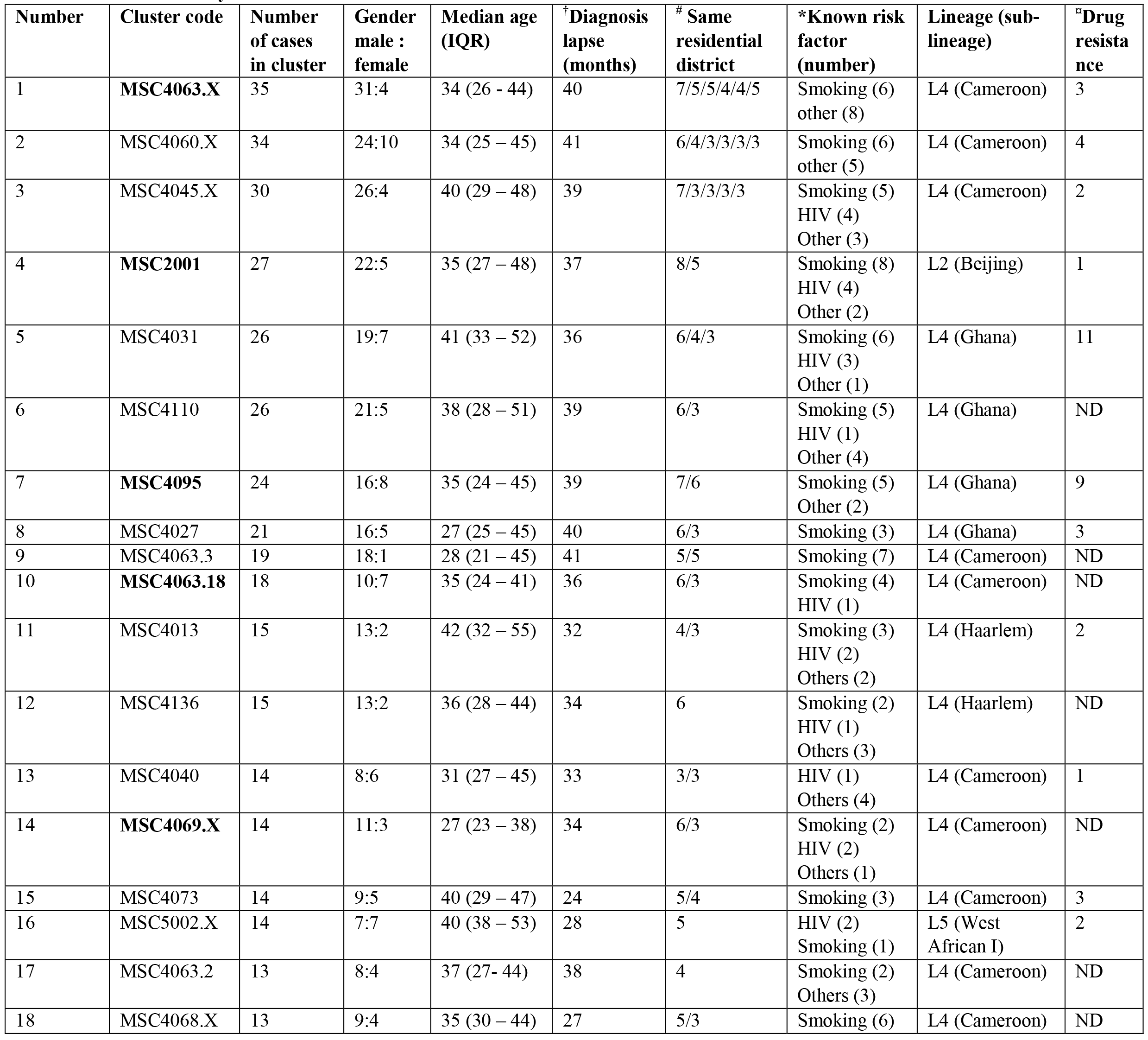
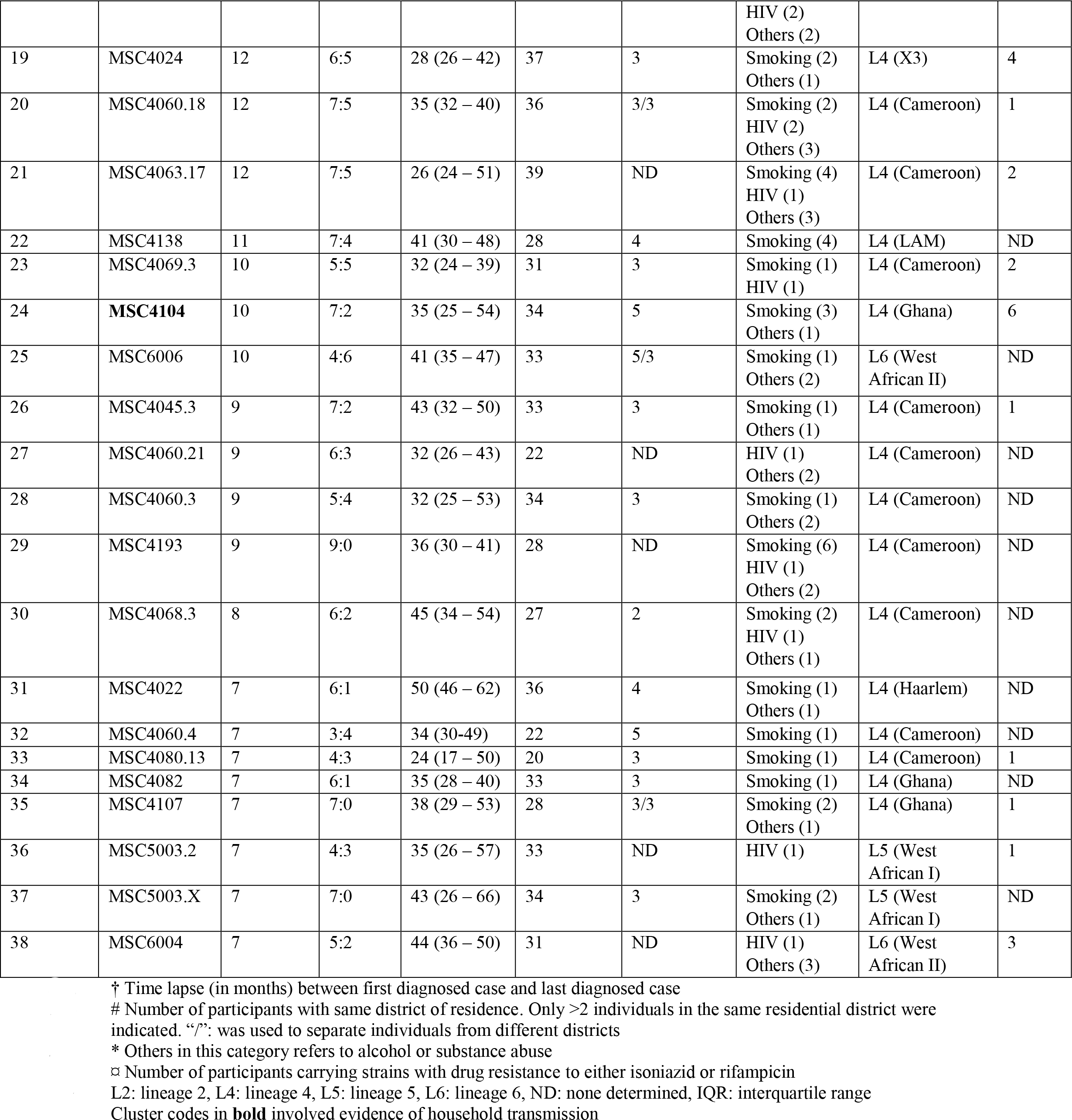
Characteristics of large molecular clusters resulting from combined 15-MIRU and spoligotyping cluster analysis

Epidemiological investigations revealed both localized and dispersed recent transmission among the clustered cases with suggested evidence of household transmission in at least six large clusters (MSC4063·X. MSC2001. MSC4095. MSC4063·18. MSC4069·X and MSC4104). Specifically. the same L4 strain (part of cluster MSC4069·X) was found to circulate among three individuals belonging to the same household with the oldest person (age 49) reporting having contact with his son who had TB four months prior to his episode (suggestive of household transmission). Majority of the large clusters involved TB strains circulating over almost the entire study period (Figure 6). Apart from three Ghana sub-lineage clusters (MSC4104. MSC4031 and MSC4095) and one L6 cluster (MSC6004) with respectively 60% (6/10). 42% (11/26). 38% (9/24) and 43% (3/7) of isolates showing resistance to either rifampicin and/or isoniazid (table 3). we did not observe such high levels of drug resistance in the other large and very large clusters. Only 2% of the isolates belonging to large and very large clusters were MDR-TB strains and this was significantly lower than that for small (2 isolates) and medium (3 − 5 isolates) (4%) clusters (p=0·031).

**Figure 6:**
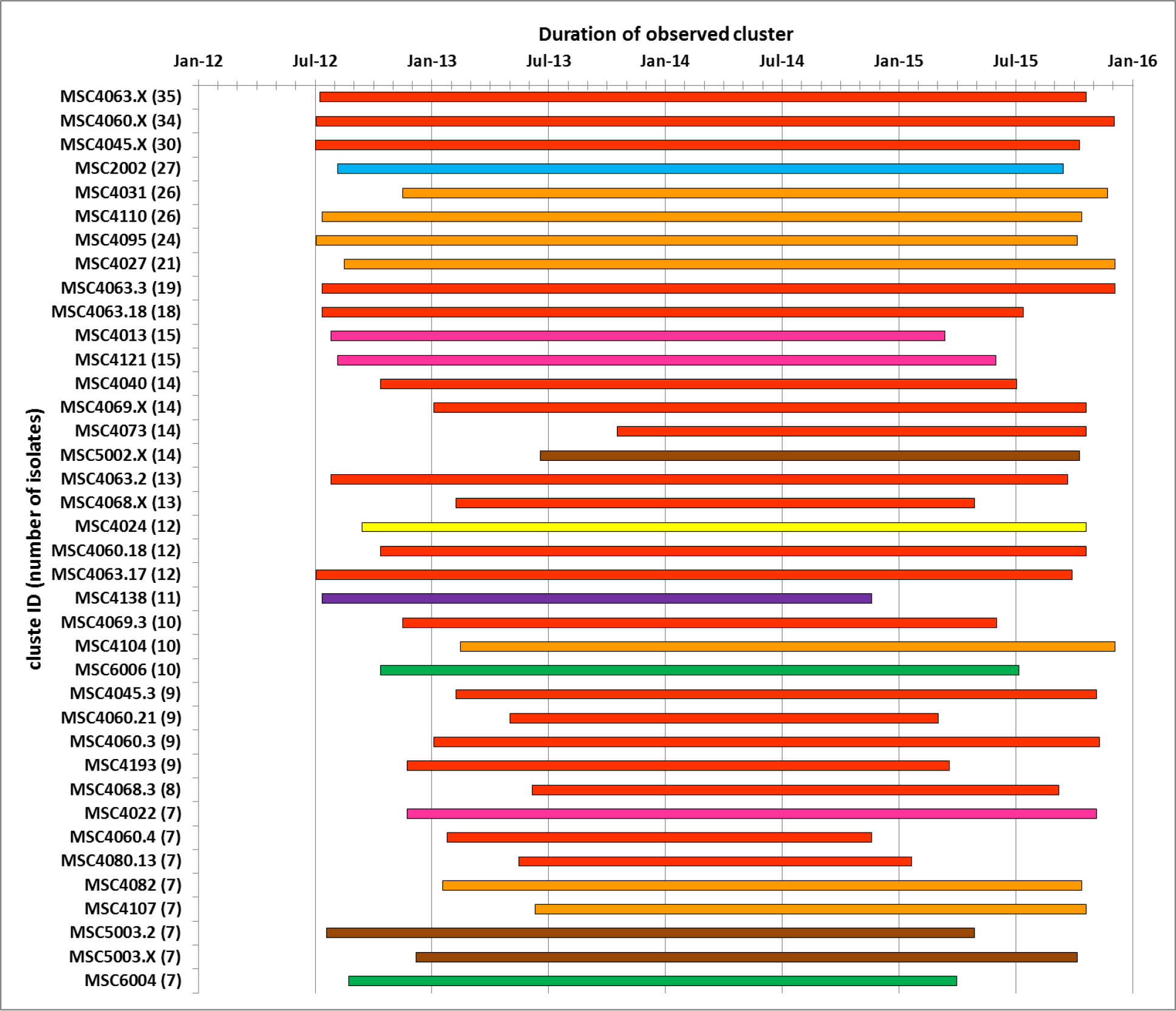
Time lapse between first and last diagnosed case within selected large and very large clusters. Lineages 5 and 6 have been color coded with the universally accepted color codes for the main MTBC lineages whereas sub-lineages of lineage 4 have been color coded, Cameroon: red, Ghana: gold, Haarlem: pink, LAM: purple and X3: yellow.

For the determination of possible factors associated with recent TB transmission. we first performed a general logistic regression model including all MTBC lineages using the event of belonging to a clustered case as the outcome variable and participants’ variables as possible predictors (table 4). In a separate logistic regression model. we tested risk factors associated with recent TB transmission stratified independently by L4 and L5 (table 5) excluding L6 due to limited sample size. In the multivariable analysis for the general logistic regression model. we found that harboring either isoniazid or rifampicin resistant MTBC strain (adjusted OR 0·7. 95% CI 0·5 − 0·9) was associated with a lower odds of belonging to a clustered case (table 4). All other factors such as education status, occupation, income level, ethnicity, religion or HIV status had no association with recent TB transmission.

Finally, using adjusted predictions, we found that, the probability of belonging to a clustered case decreased with age and increased with the number of TB contacts (Figure 7). In a separate logistic regression analysis, including age as a continuous variable with belonging to a clustered case the outcome variable, we found that, each year increase in age was significantly associated with approximately 1% (CI: 0·13 − 2·00%) decrease in the odds of a TB patient being part of a recent transmission event (p=0·007).

**Figure 7:**
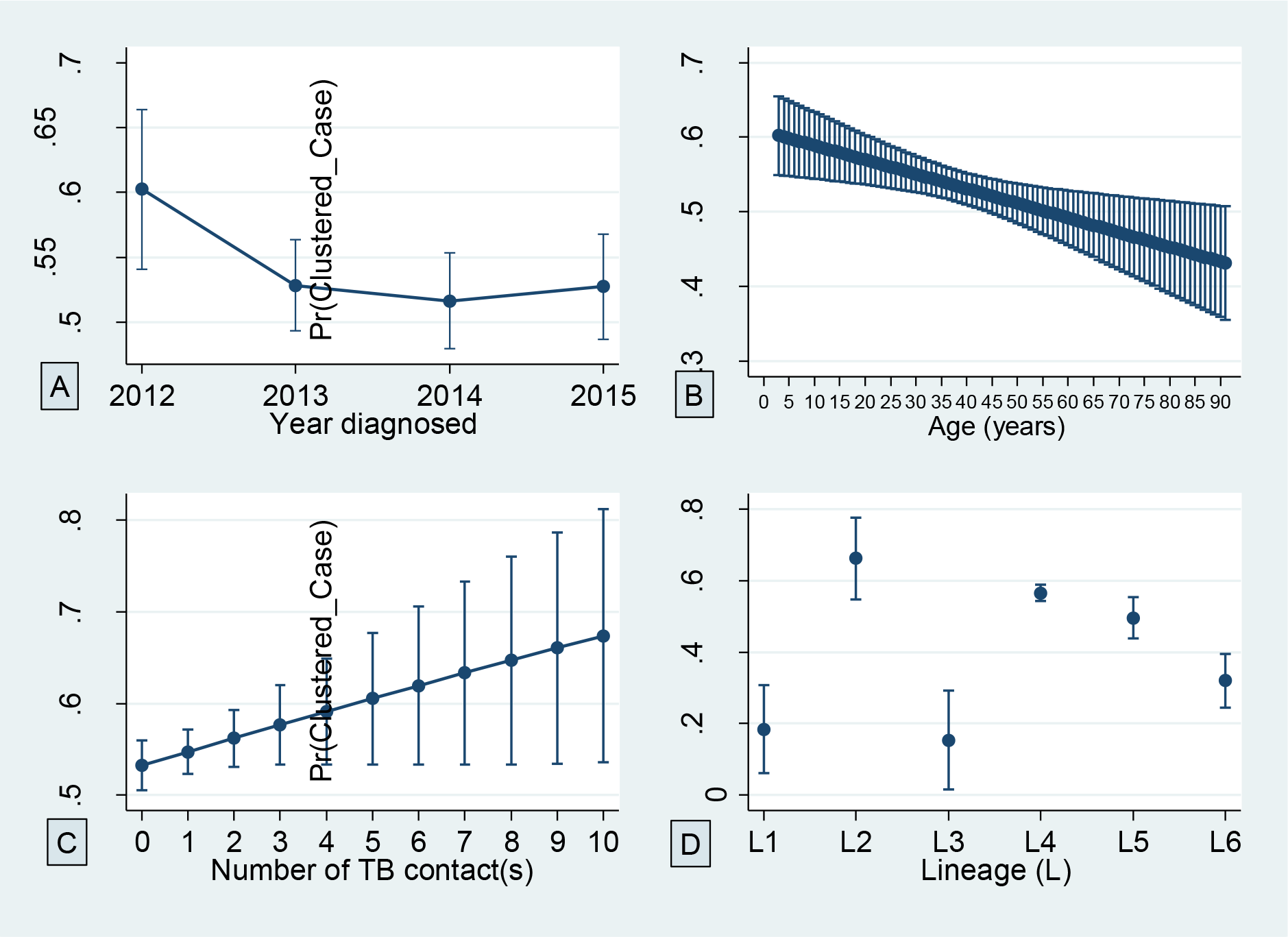
Adjusted predictions of the probability of belonging to a clustered case with 95% confidence interval at the year of diagnosis (A), while ageing (B), considering the number of close TB contact(s) (C) and of circulating MTBC lineages (D).

**Table 4:** logistic Regression analysis of risk factors associated with TB clustering (recent TB transmission)

| Variable | MTBC (N =2,309) |  | Univariate |  | Multivariate |  |
| --- | --- | --- | --- | --- | --- | --- |
|  | Total TB cases, N (%) | Clustered<sup>#</sup> cases, N (%) | OR (95% CI) | p-value | Adjusted OR (95% CI) | p-value |
| Year diagnosed | 2,309 (100) | 1,229 (53.2) |  |  |  |  |
| 2012 | 244 (10.6) | 147 (60.3) | 1.4 (1.0 - 1.8) | 0.043 | 1.3 (0.9 - 1.7) | 0.113 |
| 2013 | 776 (33.6) | 410 (52.8) | Reference |  |  |  |
| 2014 | 707 (30.6) | 365 (51.6) | 1.0 (0.8 - 1.2) | 0.642 | 0.9 (0.7 - 1.1) | 0.203 |
| 2015 | 582 (25.2) | 307(52.8) | 1.0 (0.8 - 1.2) | 0.975 | 1.0 (0.8 - 1.2) | 0.703 |
| Gender | 2,294 (99.4) |  |  |  |  |  |
| Male | 1,631 (71.1) | 863 (52.9) | 1.0 (0.8 - 1.2) | 0.685 |  |  |
| Female | 663 (28.9) | 357 (53.8) | Reference |  |  |  |
| Age<sup>‡</sup> (yrs) | 2,224 (96.3) |  |  |  |  |  |
| <15 | 37 (1.7) | 25 (67.6) | 1.6 (0.8 - 3.3) | 0.183 | 1.6 (0.8 - 3.2) | 0.221 |
| 15 - 29 | 639 (28.7) | 360 (56.3) | Reference |  |  |  |
| 30 - 39 | 570 (25.6) | 307 (53.9) | 0.9 (0.7 - 1.1) | 0.387 | 0.9 (0.7 - 1.2) | 0.688 |
| 40 - 59 | 778 (35.0) | 398 (51.2) | 0.8 (0.7 - 1.0) | 0.052 | 0.9 (0.7 - 1.1) | 0.241 |
| >59 | 200 (9.0) | 97 (48.5) | 0.7 (0.5 - 1.0) | 0.053 | 0.9 (0.6 - 1.1) | 0.211 |
| Nationality | 1,781 (77.1) |  |  |  |  |  |
| Ghanaian | 1,714 (96.2) | 932 (54.4) | Reference |  |  |  |
| Others | 67 (3.8) | 38 (56.7) | 1.1 (0.7 - 1.8) | 0.706 |  |  |
| Locality | 2,309 (100) | 1,229 (53.2) |  |  |  |  |
| Rural | 201 (8.7) | 74 (36.8) | Reference |  |  |  |
| Urban | 2,108 (91.3) | 1,155 (54.8) | 2.1 (1.5 - 2.8) | <0.001 |  |  |
| Residence classification | 1,642 (71.1) |  |  |  |  |  |
| Village | 69 (4.2) | 27 (39.1) | 0.5 (0.3 - 0.8) | 0.007 |  |  |
| Town | 182 (11.1) | 96 (52.7) | 0.9 (0.6 - 1.2) | 0.415 |  |  |
| City residential area | 52 (3.2) | 27 (51.9) | 0.8 (0.5 - 1.5) | 0.564 |  |  |
| City suburb | 1,136 (69.2) | 636 (56.0) | Reference |  |  |  |
| City Slum | 203 (12.4) | 112 (55.2) | 1.0 (0.7 - 1.3) | 0.83 |  |  |
| Residential district | 1,538 (66.6) |  |  |  |  |  |
| Ablekuma | 545 (35.4) | 298 (54.7) | Reference |  |  |  |
| Ashiedu Keteke | 170 (11.1) | 100 (58.8) | 1.2 (0.8 - 1.7) | 0.343 |  |  |
| Ayawaso | 220 (14.3) | 124 (56.4) | 1.1 (0.8 to 1.5) | 0.672 |  |  |
| Kpeshie | 224(14.6) | 121 (54.0) | 1.0 (0.7 - 1.3) | 0.867 |  |  |
| Mamprusi East | 70 (4.6) | 22 (31.4) | 0.4 (0.2 - 0.6) | <0.001 |  |  |
| Okaikoi | 176 (11.4) | 98 (55.7) | 1.0 (0.7 to 1.5) | 0.816 |  |  |
| Osu Klottey | 133 (8.6) | 78 (58.7) | 1.2 (0.8 - 1.7) | 0.409 |  |  |
| Household type | 1,624 (70.3) |  |  |  |  |  |
| Self-contain | 412 (25.4) | 221 (53.6) | 1.0 (0.8 - 1.2) | 0.797 |  |  |
| Compound house | 1,212 (74.6) | 659 (54.4) | Reference |  |  |  |
| Education | 1,748 (75.7) |  |  |  |  |  |
| Primary | 222 (12.7) | 125 (56.3) | 1.1 (0.8 - 1.5 ) | 0.637 |  |  |
| Middle/JHS | 637 (36.4) | 347 (54.5) | Reference |  |  |  |
| Secondary | 429 (24.5) | 232 (54.1) | 1.0 (0.8 - 1.3) | 0.899 |  |  |
| Tertiary | 190 (10.9) | 110 (57.9) | 1.1 (0.8 - 1.6) | 0.405 |  |  |
| No Education | 270 (15.4) | 141 (52.2) | 0.9 (0.7 - 1.2) | 0.534 |  |  |
| Occupation | 1,722 (74.6) |  |  |  |  |  |
| Unemployed | 390 (22.6) | 208 (53.3) | 0.9 (0.7 - 1.1) | 0.423 |  |  |
| Unskilled | 951 (55.2) | 530 (55.7) | Reference |  |  |  |
| Skilled | 381 (22.1) | 198 (52.0) | 0.9 (0.7 - 1.1) | 0.213 |  |  |
| Monthly income (GH¢) | 1,622 (70.2) |  |  |  |  |  |
| None | 371 (22.9) | 213 (57.4) | Reference |  |  |  |
| <301 | 807 (49.7) | 438 (54.3) | 0.9 (0.7 - 1.1) | 0.315 |  |  |
| 301 - 1,000 | 407 (25.1) | 218 (53.6) | 0.8 (0.6 - 1.1) | 0.281 |  |  |
| >1,000 | 37 (2.3) | 15 (40.5) | 0.5 (0.3 - 1.0) | 0.052 |  |  |
| Religion | 1,771 (76.7) |  |  |  |  |  |
| Christian | 1,361 (76.9) | 739 (54.3) | Reference |  |  |  |
| Islam | 302 (17.0) | 161 (53.3) | 1.0 (0.7 - 1.2) | 0.755 |  |  |
| Others | 26 (1.5) | 14 (53.9) | 1.0 (0.4 - 2.1) | 0.963 |  |  |
| Not religious | 82 (4.6) | 49 (59.7) | 1.2 (0.8 - 2.0) | 0.366 |  |  |
| Ethnicity | 1,760 (76.4) |  |  |  |  |  |
| Akan | 570 (32.3) | 309 (54.2) | Reference |  |  |  |
| Ewe | 259 (14.7) | 143 (55.2) | 1.0 (0.8 - 1.4) | 0.788 |  |  |
| Ga/Adangbe | 544 (30.8) | 310 (57.0) | 1.1 (0.9 - 1.4) | 0.352 |  |  |
| Others | 392 (22.2) | 196 (50.0) | 0.8 (0.6 - 1.1) | 0.199 |  |  |
| Marital Status | 1,758 (76.1) |  |  |  |  |  |
| Single | 766 (43.6) | 431 (56.3) | Reference |  |  |  |
| Married | 742 (42.2) | 395 (53.2) | 0.9 (0.7 - 1.1) | 0.237 |  |  |
| Divorced | 167 (9.5) | 99 (59.3) | 1.1 (0.8 - 1.6) | 0.476 |  |  |
| Widowed | 83 (4.7) | 35 (42.2) | 0.6 (0.3 - 0.9) | 0.015 |  |  |
| Smear positivity | 2,208 (95.6) |  |  |  |  |  |
| Scanty 1 - 9 | 173 (7.8) | 96 (55.5) | 1.1 (0.8 - 1.5) | 0.714 |  |  |
| 1+ | 474 (21.5) | 237 (50.0) | 0.9 (0.7 - 1.1) | 0.151 |  |  |
| 2+ | 546 (24.7) | 294 (53.9) | 1.0 (0.8 - 1.2) | 0.957 |  |  |
| 3+ | 1,015 (46.0) | 548 (54.0) | Reference |  |  |  |
| Previous TB treatment | 1,737 (75.2) |  |  |  |  |  |
| Yes | 291 (16.8) | 153 (52.6) | 0.9 (0.7 - 1.2) | 0.535 |  |  |
| No | 1,446 (83.2) | 789 (54.6) | Reference |  |  |  |
| Risk of TB contact |  |  |  |  |  |  |
| Closefriend/Household | 1,665 (72.1) |  |  |  |  |  |
| No contact | 1,121 (67.3) | 594 (53.0) | Reference |  |  |  |
| 1 contact | 212 (12.7) | 118 (55.7) | 1.1 (0.8 - 1.5) | 0.475 |  |  |
| 2-5 contacts | 309 (18.6) | 179 (57.9) | 1.2 (0.9 - 1.6) | 0.123 |  |  |
| 6-10 contacts | 23 (1.4) | 15 (65.2) | 1.7 (0.7 - 4.0) | 0.249 |  |  |
| Imprisonment | 1,660 (71.9) |  |  |  |  |  |
| Yes | 97 (5.8) | 56 (57.7) | 1.1 (0.8 - 1.7) | 0.513 |  |  |
| No | 1,563 (94.2) | 849 (54.3) | Reference |  |  |  |
| Health/Lab worker | 1,661 (71.9) |  |  |  |  |  |
| Yes | 47 (2.8) | 25 (53.2) | 0.9 (0.5 - 1.7) | 0.85 |  |  |
| No | 1,614 (97.2) | 881 (54.6) | Reference |  |  |  |
| Immunosuppressive condition | 1,695 (73.4) |  |  |  |  |  |
| Any | 893 (52.7) | 488 (54.6) | 1.0 (0.9 - 1.2) | 0.747 |  |  |
| None | 802 (47.3) | 432 (53.9) | Reference |  |  |  |
| Diabetes Mellitus | 534 (23.1) |  |  |  |  |  |
| Yes | 104 (19.5) | 54 (51.9) | 1.0 (0.7 - 1.5) | 0.957 |  |  |
| No | 430 (80.5) | 222 (51.6) | Reference |  |  |  |
| HIV status | 1,166 (50.5) |  |  |  |  |  |
| Yes | 144 (12.3) | 82 (56.9) | 1.1 (0.8 - 1.6) | 0.481 |  |  |
| No | 1,022 (87.7) | 550 (53.8) | Reference |  |  |  |
| Smoking | 1,518 (65.7) |  |  |  |  |  |
| Yes | 434 (28.6) | 237 (54.6) | 1.0 (0.8 - 1.2) | 0.949 |  |  |
| No | 1,084 (71.4) | 590 (54.4) | Reference |  |  |  |
| Substance abuse (excluding alcohol) | 1,401 (60.7) |  |  |  |  |  |
| Yes | 140 (10.0) | 84 (60.0) | 1.3 (0.9 - 1.8) | 0.172 |  |  |
| No | 1,261 (90.0) | 680 (53.9) | Reference |  |  |  |
| Substance abuse (including alcohol) | 1,474 (63.8) |  |  |  |  |  |
| Yes | 460 (31.2) | 250 (54.3) | 1.0 (0.8 - 1.3) | 0.858 |  |  |
| No | 1,014 (68.8) | 546 (53.8) | Reference |  |  |  |
| Lineage | 2,309 (100) |  |  |  |  |  |
| Lineage 1 | 38 (1.7) | 7 (18.4) | 0.2 (0.08 - 0.4) | <0.001 | 0.13 (0.05 - 0.36) | <0.001 |
| Lineage 2 | 65 (2.8) | 43 (66.2) | 1.5 (0.9 - 2.5) | 0.126 | 1.5 (0.9 - 2.5) | 0.155 |
| Lineage 3 | 26 (1.1) | 4 (15.4) | 0.1 (0.05 - 0.4) | <0.001 | 0.15 (0.05 - 0.45) | 0.001 |
| Lineage 4 | 1,741 (75.4) | 984 (56.5) | Reference |  |  |  |
| Lineage 5 | 289 (12.5) | 143 (49.5) | 0.8 (0.6 - 1.0) | 0.026 | 0.7 (0.6 - 0.9) | 0.032 |
| Lineage 6 | 150 (6.5) | 48 (32.0) | 0.4 (0.3 - 0.5) | <0.001 | 0.3 (0.2 - 0.5) | <0.001 |
| Lineage 4 sub-lineage |  |  |  |  |  |  |
| Cameroon | 1,046 (60.1) | 616 (58.9) | Reference |  |  |  |
| Ghana | 376 (21.6) | 206 (54.8) | 0.8 (0.7 - 1.1) | 0.167 |  |  |
| Harleem | 156 (9.0) | 91(58.3) | 1.0 (0.7 - 1.4) | 0.895 |  |  |
| LAM | 50 (2.9) | 25 (50.0) | 0.7 (0.4 - 1.2) | 0.215 |  |  |
| Uganda | 40 (2.3) | 16 (40.0) | 0.5 (0.2 - 0.9) | 0.02 |  |  |
| Others | 51 (2.9) | 26 (51.0) | 0.7 (0.4 - 1.3) | 0.265 |  |  |
| Not determined | 22 (1.3) | 4 (18.2) | 0.2 (0.1 - 0.5) | 0.001 |  |  |
| Drug resistance | 2,300 (99.6) |  |  |  |  |  |
| Any | 313 (13.6) | 138 (44.1) | 0.6 (0.5 - 0.8) | <0.001 | 0.7 (0.5 - 0.9) | 0.002 |
| None | 1,987 (86.4) | 1,090 (54.9) | Reference |  |  |  |
| Isoniazid mono-resistant | 2,300 (99.6) |  |  |  |  |  |
| Yes | 295 (12.8) | 129 (43.7) | 0.6 (0.5 - 0.8) | <0.001 |  |  |
| No | 2,005 (87.2) | 1,099 (54.8) | Reference |  |  |  |
| Multidrug resistant (MDR) | 2,300 (99.6) |  |  |  |  |  |
| Yes | 81 (3.5) | 35 (43.2) | 0.7 (0.4 - 1.0) | 0.063 |  |  |
| No | 2,219 (96.5) | 1,193 (53.8) | Reference |  |  |  |
| Cluster size (n) | 1,227 (53.1) |  |  |  |  |  |
| Small (2) | 290 (23.6) |  |  |  |  |  |
| Medium (3 - 5) | 262 (21.4) |  |  |  |  |  |
| Large (6 - 20) | 452 (36.8) |  |  |  |  |  |
| Very large (>20) | 223 (18.2) |  |  |  |  |  |

For the multivariate model, we included only variables with $p < 0.1$ and with at least 90% of available data. We however excluded "locality" due to reduced sample size from the rural setting.
We excluded residence classification, marital status, isoniazid mono-resistance and MDR due to colinearity with other variables in the model.
\* $p < 0.05$ ; \*\* $p < 0.001$

**Table 5:** Risk Factors associated with TB clustering: logistic Regression Analysis stratified by lineage

| Variables | lineage 4 (N=1,741) |  | Univariate | Multivariate |  | lineage 5 (N=289) |  | Univariate |  |
| --- | --- | --- | --- | --- | --- | --- | --- | --- | --- |
|  | TB cases, N (%) | clustered<sup>#</sup> cases, N (%) | OR (95% CI) | Adjusted OR (95% CI) | p-value | TB cases, N (%) | clustered<sup>#</sup> cases, N (%) | OR (95% CI) | p-value |
| Year diagnosed | 1,741 (100) |  |  |  |  | 289 (100) |  |  |  |
| 2012 | 183 (10.5) | 120 (65.6) | 1.5 (1.1 - 2.1)* | 1.4 (1.0 - 2.1) | 0.062 | 26 (9.0) | 14 (53.8) | 1.2 (0.5 - 2.9) | 0.659 |
| 2013 | 568 (32.6) | 318 (56.0) | Reference |  |  | 98 (33.9) | 48 (49.0) | Reference |  |
| 2014 | 548 (31.5) | 300 (54.7) | 1.0 (0.8 - 1.2) | 1.0 (0.7 - 1.3) | 0.847 | 92 (31.8) | 43 (46.7) | 0.9 (0.5 - 1.6) | 0.757 |
| 2015 | 442 (25.4) | 244 (55.2) | 1.0 (0.8 - 1.2) | 1.0 (0.7 - 1.3) | 0.955 | 73 (25.3) | 38 (52.1) | 1.2 (0.6 - 2.1) | 0.691 |
| Age (yrs) | 1,672 |  |  |  |  | 283 |  |  |  |
| <15 | 27 (1.6) | 20 (74.1) | 2.1 (0.9 - 5.0) |  |  | 5 (1.8) | 3 (60.0) |  |  |
| 15 - 29 | 497 (29.7) | 289 (58.2) | Reference |  |  | 78 (27.6) | 42 (53.8) |  |  |
| 30 - 39 | 432 (25.8) | 252 (58.3) | 1.0 (0.8 - 1.3) |  |  | 68 (24.0) | 31 (45.6) |  |  |
| 40 - 59 | 580 (34.7) | 315 (54.3) | 0.9 (0.7 - 1.1) |  |  | 94 (33.2) | 48 (51.1) |  |  |
| >59 | 136 (8.1) | 71 (52.2) | 0.8 (0.5 - 1.2) |  |  | 38 (13.4) | 16 (42.1) |  |  |
| Locality | 1,741 (100) |  |  |  |  | 289 (100) |  |  |  |
| Rural | 153 (8.8) | 59 (38.6) | Reference |  |  | 15 (5.2) | 4 (26.7) | Reference |  |
| Urban | 1,588 (91.2) | 923 (58.1) | 2.2 (1.6 - 3.1)** |  |  | 274 (94.8) | 139 (50.7) | 2.8 (0.9 - 9.1) | 0.081 |
| Residential district | 1,165 |  |  |  |  | 189 |  |  |  |
| Ablekuma | 412 (35.4) | 237 (57.5) | Reference |  |  | 77 (40.7) | 39 (50.7) | Reference |  |
| Ashiedu Keteke | 132 (11.3) | 81 (61.4) | 1.2 (0.8 - 1.8) |  |  | 13 (6.9) | 5 (38.5) | 0.6 (0.2 - 2.0) | 0.419 |
| Ayawaso | 178 (15.3) | 111 (62.4) | 1.2 (0.8 - 1.8) |  |  | 21 (11.1) | 7 (33.3) | 0.5 (0.2 - 1.4) | 0.163 |
| Kpeshie | 166 (14.2) | 88 (53.0) | 0.8 (0.6 - 1.2) |  |  | 37 (19.6) | 25 (67.6) | 2.0 (0.9 - 4.6) | 0.091 |
| Mamprusi East | 56 (4.8) | 19 (33.9) | 0.4 (0.2 - 0.7)* |  |  | 4 (2.1) | 1 (25.0) | 0.32 (0.03 - 3.26) | 0.339 |
| Okaikoi | 134 (11.5) | 80 (59.7) | 1.1 (0.7 - 1.6) |  |  | 24 (12.7) | 12 (50.0) | 1.0 (0.4 - 2.4) | 0.956 |
| Osu Klottey | 87 (7.5) | 58 (66.7) | 1.5 (0.9 - 2.4) |  |  | 13 (6.9) | 5 (38.5) | 0.6 (0.2 - 2.0) | 0.419 |
| Monthly income (GH¢) | 1,222 |  |  |  |  |  |  |  |  |
| None | 275 (22.5) | 167 (60.7) | Reference |  |  |  |  |  |  |
| <301 | 605 (49.5) | 351 (58.0) | 0.9 (0.7 - 1.2) |  |  |  |  |  |  |
| 301 - 1,000 | 314 (25.7) | 184 (58.6) | 0.9 (0.7 - 1.3) |  |  |  |  |  |  |
| >1,000 | 28 (2.3) | 11 (39.3) | 0.4 (0.2 - 0.9)* |  |  |  |  |  |  |
| Marital Status | 1,322 |  |  |  |  |  |  |  |  |
| Single | 591 (44.7) | 355 (60.1) | Reference |  |  |  |  |  |  |
| Married | 549 (41.5) | 312 (56.8) | 0.9 (0.7 - 1.1) | 0.9 (0.7 - 1.2) | 0.589 |  |  |  |  |
| Divorced | 124 (9.4) | 78 (62.9) | 1.1 (0.8 - 1.7) | 1.1 (0.7 - 1.7) | 0.543 |  |  |  |  |
| Widowed | 58 (4.4) | 24 (41.4) | 0.5 (0.3 - 0.8)* | 0.5 (0.3 - 0.8) | 0.011 |  |  |  |  |
| Lineage 4 sub-lineage |  |  |  |  |  |  |  |  |  |
| Cameroon | 1046 (60.1) | 614 (58.7) |  |  |  |  |  |  |  |
| Ghana | 376 (21.6) | 206 (54.8) | 0.9 (0.7 - 1.1) | 0.9 (0.7 - 1.2) | 0.403 |  |  |  |  |
| Harleem | 156 (9.0) | 91 (58.3) | 1.0 (0.7 - 1.4) | 1.0 (0.7 - 1.5) | 0.87 |  |  |  |  |
| LAM | 50 (2.9) | 25 (50.0) | 0.7 (0.4 - 1.2) | 0.7 (0.4 - 1.4) | 0.354 |  |  |  |  |
| Uganda | 40 (2.3) | 16 (40.0) | 0.5 (0.2 - 0.9)* | 0.4 (0.2 - 0.8) | 0.013 |  |  |  |  |
| Others | 51 (2.9) | 26 (51.0) | 0.7 (0.4 - 1.3) | 0.8 (0.4 - 1.6) | 0.558 |  |  |  |  |
| Not determined | 22 (1.3) | 4 (18.2) | 0.2 (0.1 - 0.5)* | 0.10 (0.03 - 0.35) | <0.001 |  |  |  |  |
| Drug resistance | 1,736 |  |  |  |  |  |  |  |  |
| Any | 241 (13.9) | 114 (47.3) | 0.7 (0.5 - 0.9)* | 0.7 (0.5 - 1.0) | 0.059 |  |  |  |  |
| None | 1,495 (86.1) | 867 (58.0) | Reference |  |  |  |  |  |  |

We included in this analysis, only variables with $p < 0.1$ from the general logistic regression model in table 4
For the multivariate model, we included only variables with $p < 0.1$ and with at least 90% of available data.
\* $p < 0.05$ ; \*\* $p < 0.001$

## Discussion

The aims of this study were to conduct a population-based prospective molecular epidemiological study to analyze the transmission dynamics of MTBC strains circulating in Ghana and to identify risk factors associated with recent TB transmission. The current study also offered us the opportunity to create a local repository of strain types for future reference to help monitor TB programs by analyzing the trends in estimates of recent transmission.

We obtained a high MTBC isolate recovery rate of 78·8% higher than that reported in similar studies^28,29^ and this strenghtens the power of the sample size to make assessments of TB transmission rate in Ghana. Our study identified a high TB clustering (recent TB transmission) rate of 41·2% which is quite alarming with the urban and rural areas having estimated rates of 41·7% and 9·0% respectively (table 2b). These findings call for intensifying community outreach programs to encourage early case reporting and infection control. Moreover, our analysis predicted the probability of clustering to generally increase with increase in the number of TB contacts (Figure 7). This means that a susceptible individual is likely to have TB and be involved in a recently transmitted event as the number of TB contacts increases.

Within our study population, we found no association of recent TB transmission with education status, occupation, income level, ethnicity, religion or HIV status. However, we observed that individuals below the age of 30 years were associated with recent TB transmission, and this is similar to observations made elsewhere. ^28,30^ Also from this study, we observed that each year increase in age is associated with an approximately 1% (CI: 0·13 − 2·00, p=0·007) decrease in the odds of a TB patient being part of a recent transmission event implying that, compared to younger individuals, older individuals are more likely to get active TB disease by reactivation of latent TB infection rather than through a recent transmission event.^28^ This finding put age as a risk factor for recent TB transmission in Ghana. Also, we identified that the male/female ratio among very large clusters was significantly higher than that observed in the general TB patient population (p=0·022). This finding, together with the observation that some large clusters involved only males, also puts males as having a higher risk of recent TB transmission compared to females suggesting that males may engage in certain social activities that predisposes them to belonging to a recent transmission event.

We observed a lower rate of MDR-TB among large clustered cases compared to the general population (2% vs 4%, p=0·031) indicating a low MDR-TB transmissibility within our study population. This finding further suggests that, majority of drug resistant-TB cases in Ghana acquired the drug resistance during treatment which indicate poor patient compliance.^31^ We also found that compared to drug (isoniazid and/or rifampicin) sensitive TB strains, it is unlikely to find TB strains with either isoniazid and/or rifampicin resistance involved in a recent transmission event (adjusted OR 0·7, 95% CI 0·5 − 0·9).

Within our study setting, we observed a reduced transmission of Maf (L5: 31·8%, L6: 24·7%) compared to MTBss L4 (44·9%). The high recent transmission rate observed for L4 was driven by both the Cameroon and Ghana sub-lineages with no difference in their transmissibility, hence putting the Ghana sub-lineage as a very important pathogen. The high transmission of the Ghana sub-lineage coupled with recently reported association with drug resistance^18^ is of public health importance and hence calls for the national tuberculosis control program to support peripheral diagnostic laboratories with facilities to accurately detect and help control the spread of the Ghana sub-lineage.

The higher recent transmission rate for L4 compared to L5 may arguably just be a mirage and may not necessarily imply the outcompeting of L5 by L4 as their relative proportions still remain constant over the entire study period of this current study (Figure 3) and also from previous reports.^16^ The reduced prevalence and transmission of L5 may just be a result of a bottleneck event which might have taken place in the past resulting in the reduction of Maf population at the time and ever since. Maf has been transmitting at this same average rate ensuring a stable population proportion over time. Despite the low transmissibility of Maf. the observed stable relative proportion over the entire study period may be because the pathogen. has adapted to infecting specific host populations (possibly due to unidentified host genetic or environmental factors peculiar to some West African inhabitants) and hence enabling the maintenance of a stable prevalence over time. Using adjusted predictions for the probability of clustering. we found that. Maf L5 may still have the propensity to transmit as equal as lineage 4 (Figure 7) not forgetting the confounding effect of a higher diversity in spoligotype pattern of L5 compared to o L4 and hence reducing the clustering of the former.^8^ Compared to L4. we found a significant association of L6 with individuals living in villages (OR 6·6. p<0·05; appendix p 3). The low recent TB transmission in the villages coupled with an association of L6 could be the reason why we observed low frequencies of L6 strains within our study setting.

Our report could be limited by the possibility of underestimating recent transmission rate resulting from misclassifying strains as unique if they were actually clustered outside of our restricted geographic sampling site and sampling period. We however took some measures to address underestimation of recent TB transmission by recruiting up to 90% of the diagnosed TB cases spanning a three and a half year period. In addition. we also understand the possibility of overestimating recent TB transmission rates considering that the basis of our clustering analysis was done using a combined 15 loci MIRU-VNTR typing and spoligotyping whereas WGS could have offered a better resolution of strains.

Overall, our findings indicate high recent TB transmission suggesting occurrences of unsuspected outbreaks and recommends intensifying community education to improve early case reporting and infection control.

## Contributors

DYM designed the study, provided supervision and support, provided intellectual input and wrote the manuscript. PA performed most of the laboratory procedures, collated the epidemiological and laboratory data, did all statistical and cluster analysis on the data and wrote the manuscript. AAP contributed to the study design, performed some laboratory procedures and provided all associated data and provided useful comments to writing the manuscript. DAP performed some laboratory procedures and provided all associated data. SB provided supervision, support, intellectual input and critically reviewed the manuscript. SOW performed some laboratory procedures and provided all associated data and provided useful comments to writing the manuscript. IDO performed some laboratory procedures and provided all associated data and provided useful comments to writing the manuscript. AF supported the enrollment and collection of clinical and demographic data from the health facilities. GA did some laboratory procedures and provided all associated data. KAK provided supervision and support, intellectual input and provided useful comments to writing the manuscript. SG designed the study, provided supervision and support, provided intellectual input and critically reviewed the manuscript. All coauthors reviewed and approved the final manuscript before submission.

## Acknowledgements

The research was funded with the Wellcome Trust Intermediate Fellowship Grant 097134/Z/11/Z to Dorothy Yeboah-Manu. PA was supported by the West African Centre for Cell Biology of Infectious Pathogens (WACCBIP)-World Bank ACE PhD studentship. The authors are grateful for the administrative support of Dr. Frank Bonsu, National Tuberculosis Control Program, Ghana, laboratory heads and nurses at various facilities we recruited cases and all study participants. We thank the national service personnel for providing immense help in completing questionnaires and making sputum samples available for laboratory investigations. We thank Vida Yirenkyiwaa Adjei and Portia Abena Morgan of Noguchi Memorial Institute for Medical Research for their assistance in some laboratory procedures.

